# Single shot detector application for image disease localization

**DOI:** 10.1101/2021.09.21.461307

**Authors:** Rushikesh Chopade, Aditya Stanam, Shrikant Pawar

## Abstract

Bounding box algorithms are useful in localization of image patterns. Recently, utilization of convolutional neural networks on X-ray images has proven a promising disease prediction technique. However, pattern localization over prediction has always been a challenging task with inconsistent coordinates, sizes, resolution and capture positions of an image. Several model architectures like Fast R-CNN, Faster R-CNN, Histogram of Oriented Gradients (HOG), You only look once (YOLO), Region-based Convolutional Neural Networks (R-CNN), Region-based Fully Convolutional Networks (R-FCN), Single Shot Detector (SSD), etc. are used for object detection and localization in modern-day computer vision applications. SSD and region-based detectors like Fast R-CNN or Faster R-CNN are very similar in design and implementation, but SSD have shown to work efficiently with larger frames per second (FPS) and lower resolution images. In this article, we present a unique approach of SSD with a VGG-16 network as a backbone for feature detection of bounding box algorithm to predict the location of an anomaly within chest X-ray image.

## Introduction

Object localization is a subfield of computer vision that is used to detect the location of object in an image. Several model architectures like Fast R-CNN [1], Faster R-CNN [2], Histogram of Oriented Gradients (HOG) [3], You only look once (YOLO) [4], Region-based Convolutional Neural Networks (R-CNN) [5], Region-based Fully Convolutional Networks (R-FCN) [6], Single Shot Detector (SSD) [7] and Spatial Pyramid Pooling (SSP-net) [8] are been used for object detection and localization in modern-day computer vision applications. The SSD and region-based detectors like Fast R-CNN or Faster R-CNN are very similar in design and implementation, but SSD have shown to work efficiently with larger frames per second (FPS) and lower resolution images [7]. Although Region-based detectors like Faster R-CNN have a little greater accuracy as compared to SSD, SSD’s are faster and better for real-time image processing [9]. Thus, we present a unique approach of SSD with a VGG-16 network as a backbone for feature detection of bounding box algorithm to predict the location of an anomaly within chest X-ray image.

## Method

### 1) Data collection

The image dataset for developing bounding box algorithms has been retrieved from National Institutes of Health (NIH) kaggle portal. The dataset consists of 112,120 chest X-ray images, each image with a 1024^*^1024-pixel resolution. The images are divided into 15 classes (‘No Finding’, ‘Atelectasis’, ‘Cardiomegaly’, ‘Consolidation’, ‘Effusion’; ‘Emphysema’, ‘Edema’, ‘Fibrosis’, ‘Infiltration’, ‘Mass’, ‘Nodule’, ‘Pneumonia’, ‘Pneumothorax’, ‘Pleural Thickening’ and ‘Hernia’). Further, each X-ray image consists of information on 4 bounding box attributes which bound the exact location of the detected disease. The first coordinate (x_min) marks the x coordinate of the top left corner of the bounding box which can be considered as the origin with pixels measured from this corner of the image. Similarly, the second attribute (y_min) marks the y coordinate of the top left corner of the bounding box. The remaining two attributes are the width and height of the bounding box in unit pixels length. Figure 1 is the X-ray image of a patient suffering from cardiomegaly. The red bounding box shows the location of the infection in the image. The image is downscaled to 512^*^512 from 1024^*^1024, 1024^*^1024 being the original resolution of X-ray image. The top left corner of the image (0,0) is taken as the origin with x_min and y_min as the x and y coordinates of bounding boxes.

**Figure 1:**
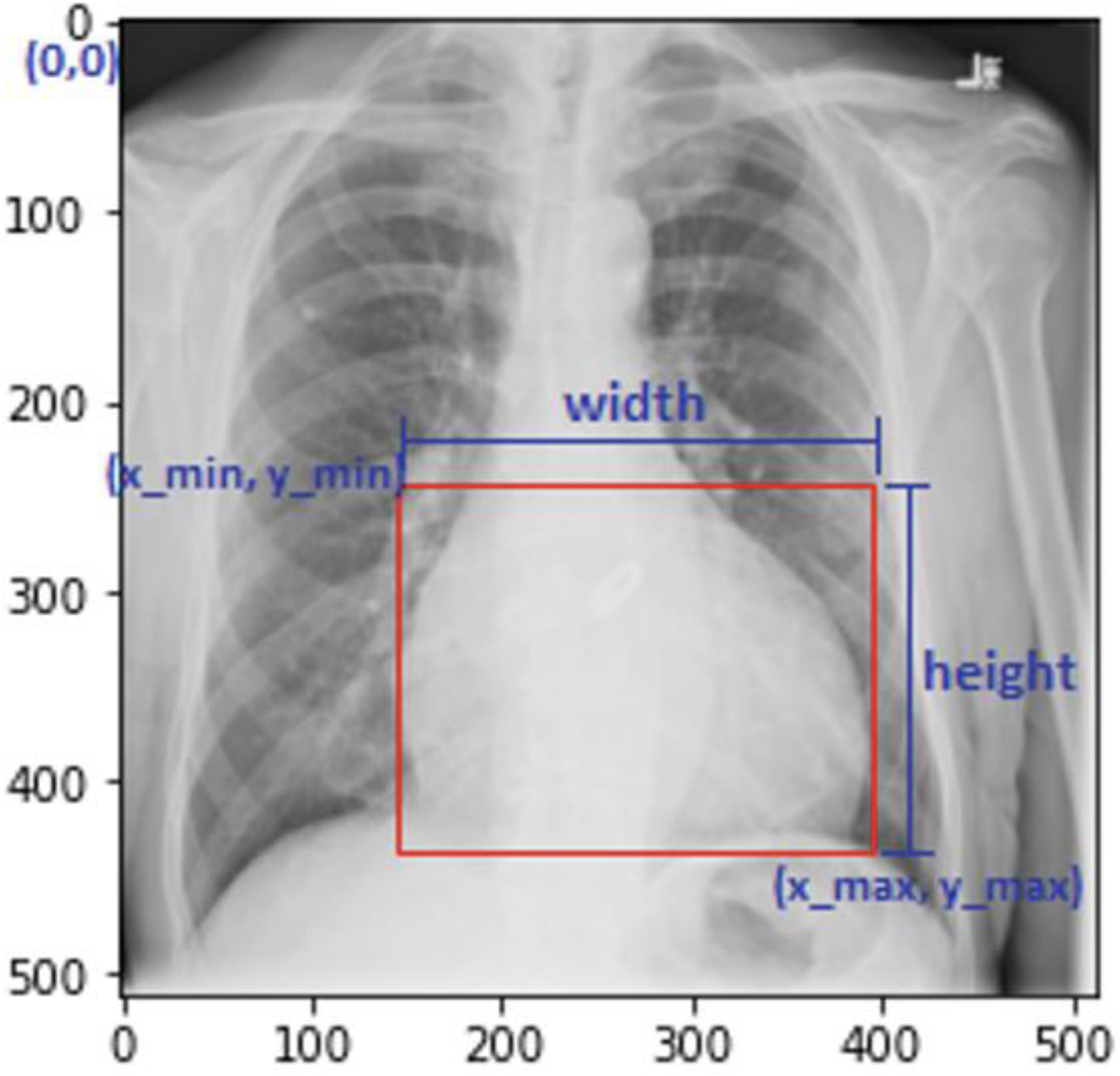
Depicts the location of cardiomegaly with bounding boxes.

### 2) Exploratory data analysis and preprocessing

For training, the width and height attributes were converted to x_max and y_max by adding the width attribute of the bounding box to the corresponding x_min coordinate for obtaining x_max and by adding the height attribute to the y_min for obtaining y_max coordinate. Thus, the bounding box coordinates can now be presented with a string containing x_min, y_min, x_max, and y_max coordinates. The images with multiple labels (93) would create a situation of high bias and abide the algorithm from learning the location of the disease with precision. Therefore, the images with multiple labels have not been included in training the algorithms. In total, 787 images with single labels have only been considered for training. The plot for the top 15 labels (single + multiple) has been shown in figure 2.

**Figure 2:**
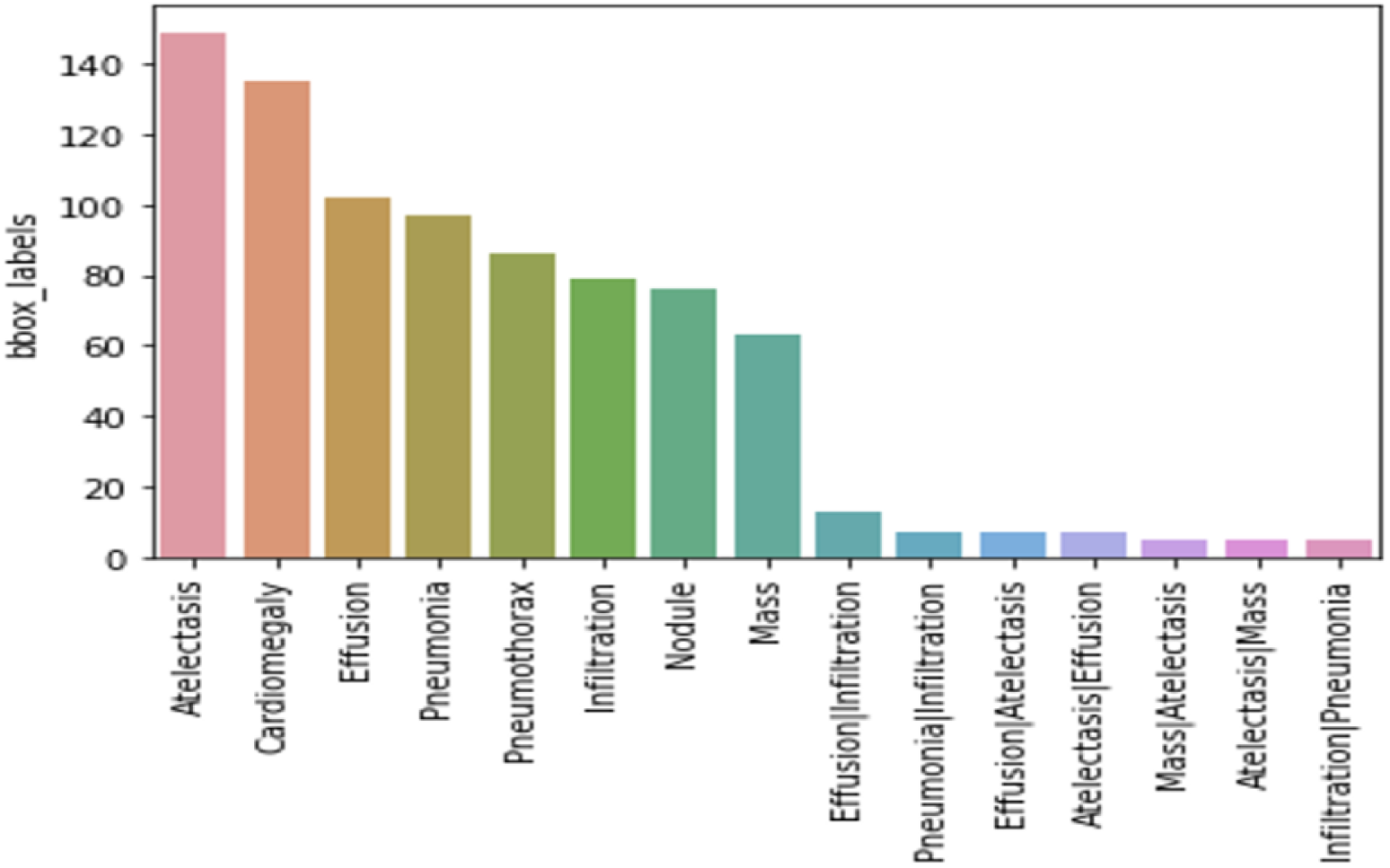
The plot for the top 15 labels.

Dynamic training has been implemented to reduce the computational cost with weights updated by backpropagation for every 4 images. An image data generator class has been utilized for this technique.

**Table 1:**
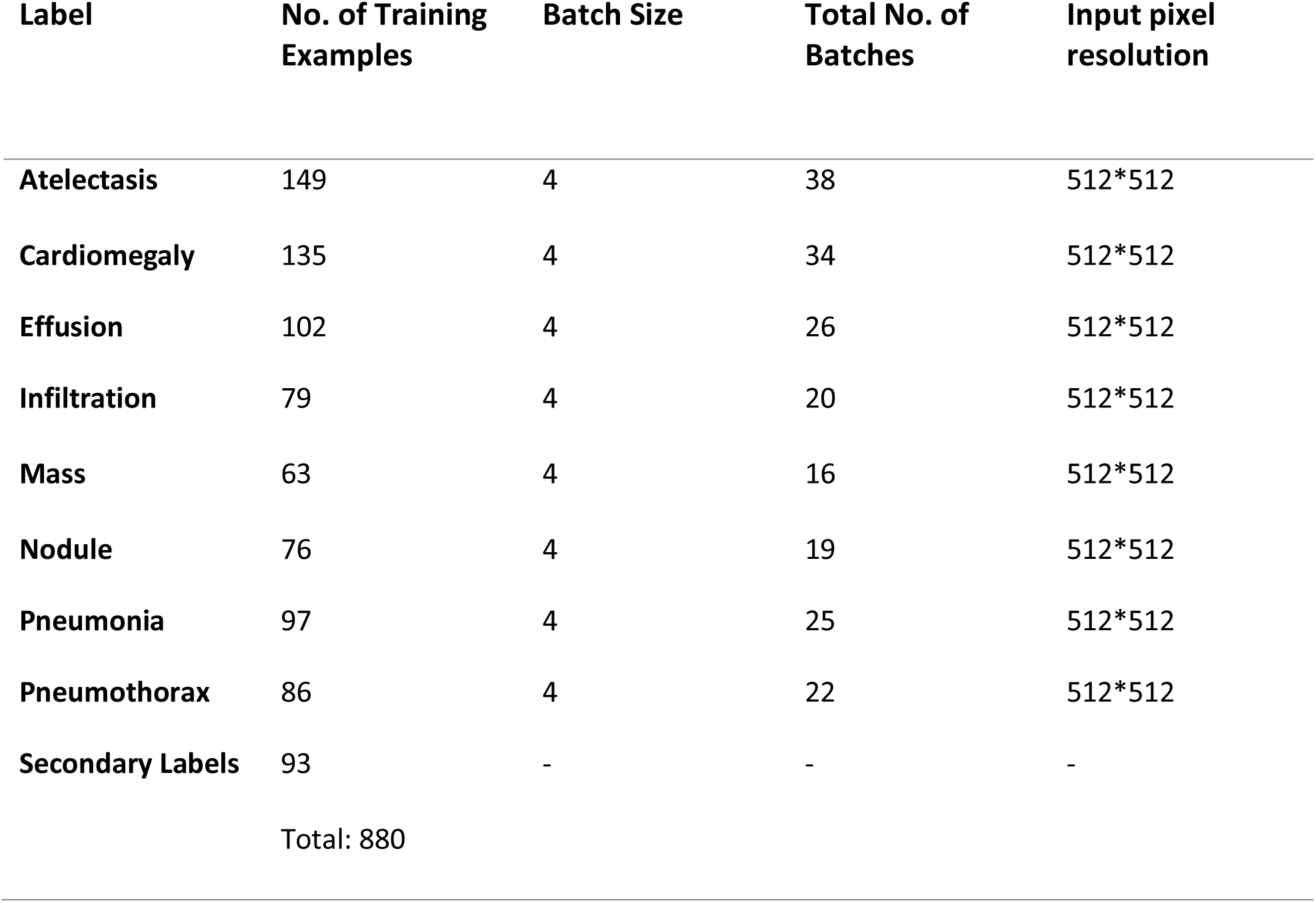
Number of training batches, sizes and input pixel resolution for bounding box algorithms.

### 3) Network architecture

Several factors can impact the accuracy and training speed of algorithm, some can be feature extractors (VGG-16, InceptionNet, ResNet, MobileNet, etc.), input image resolutions, matching strategy and IOU threshold, non-max suppression IOU threshold, number of predictions, boundry box encoding, data augmentation, size of training dataset, use of multi-scale images in training and testing, training configurations including batch size, input image resize, learning rate, learning rate decay and localization loss function [9]. An SSD runs a convolutional network on input image and computes a feature map, it then runs n^*^n convolutional kernels on this feature map to predict the bounding boxes and categorization probability [10]. For this SSD model, a VGG-16 feature extractor has been used with pretrained ImageNet weights with an input size of 512^*^512^*^3 resolution. The reason for using a feature extractor was to get the features of objects in specific order, which eventually would help the algorithm learn faster. VGG-16 feature extractor is followed by rectified linear activation layer, followed by a dropout layer to construct algorithms backbone. A dropout layer with 25% dropout nodes has been used to address the high variance problem. The output image after the dropout layer application has a dimension of 16^*^16^*^512 resolution. After the dropout layer, compression layers are added to the model architecture, containing a 2D convolutional layer, a ReLU activation layer, and a batch normalization layer. The detailed structure of this compression layer is shown in figure 3. The first compression layer has a convolutional layer of kernel size 3, a stride of 1, number of filters as 256. The output shape is obtained by the following formula:

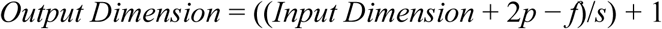

**Figure 3:**
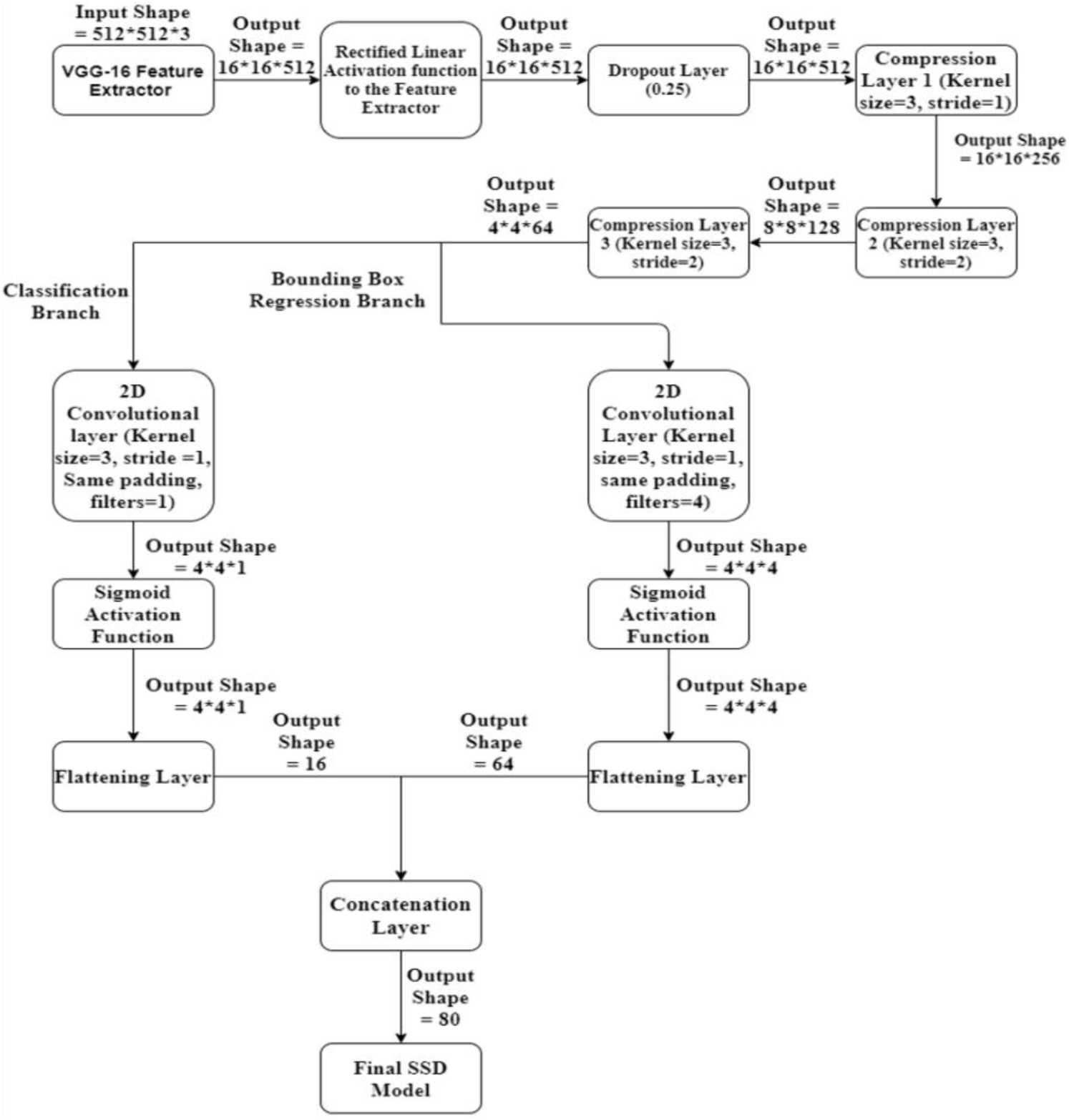
The SSD model architecture.

Where, “*p*” is padding, “*f*” is kernel size, and “*s*” is the stride used in layer. For convolutional layer of the first compression layer, with p=1, f=3, and s=1 generates an output shape 16^*^16^*^256 resolution. The second compression layer with p=1, f=3, s=1 and number of filters=128, generates an output shape of 8^*^8^*^128, this is followed by the last compression layer. After 3 compression layers, the model splits into 2 branches, classification branch and a bonding box regression branch. The classification branch classifies the image into the classes from the given labels. The 2D convolutional layer is followed by an activation layer containing the sigmoid activation function, followed by a flattening layer. The final output shape of the classification breach after flattening is equal to 16. With similar approach on bounding box regression branch, the final output shape of the classification breach after flattening equals to 64. The flattened layers of the classification and bounding box regression branches are concatenated to get final output shape with 16 bounding box predictions. The non-max suppression technique is applied to generate a single confidence value.

### 4) Custom cost function

The compression layer has been designed to increase the number of filters/channels. The 2D convolutional layer is followed by a rectified linear (ReLU) activation function. The ReLU is followed by a Batch normalization layer, which helps to stabilize the learning process and dramatically reduces the number of training epochs required to train the network. A 0.25 fraction dropout regularization has been applied to the model in order to reduce the degree of overfitting, 25% of the random nodes have been dropped with remaining nodes as input for the next hidden layer. The final SSD model after concatenation requires an additional custom loss function for training.

A custom cost function has been used to minimize the loss while training SSD algorithm. The cost function has two parts. One-part deals with classification loss while the other part deals with bounding box loss. This bounding box loss part of the custom cost function is based on intersection over union policy. In figure 5, the predicted bounding box of a patient suffering from cardiomegaly is shown in blue color. The original bounding box presenting the ground truth is shown in red color. The region of intersection of the two bounding boxes is shown in green color. The union of the two bounding boxes is simply the total area occupied by both the bounding boxes. The intersection over union is defined as area of region of intersection divided by the area of the region of union. The more is the intersection over union for the image, the lesser is the training loss. This custom cost function works to decrease the loss by increasing the area of overlap for two bounding boxes.

**Figure 4:**
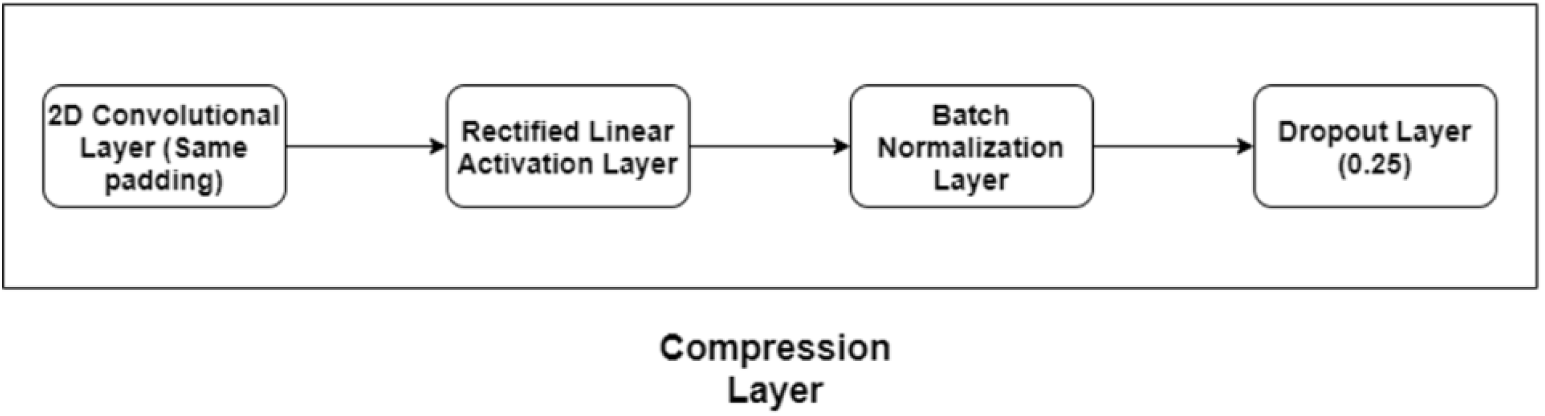
Compression layer architecture used in SSD model.

**Figure 5:**
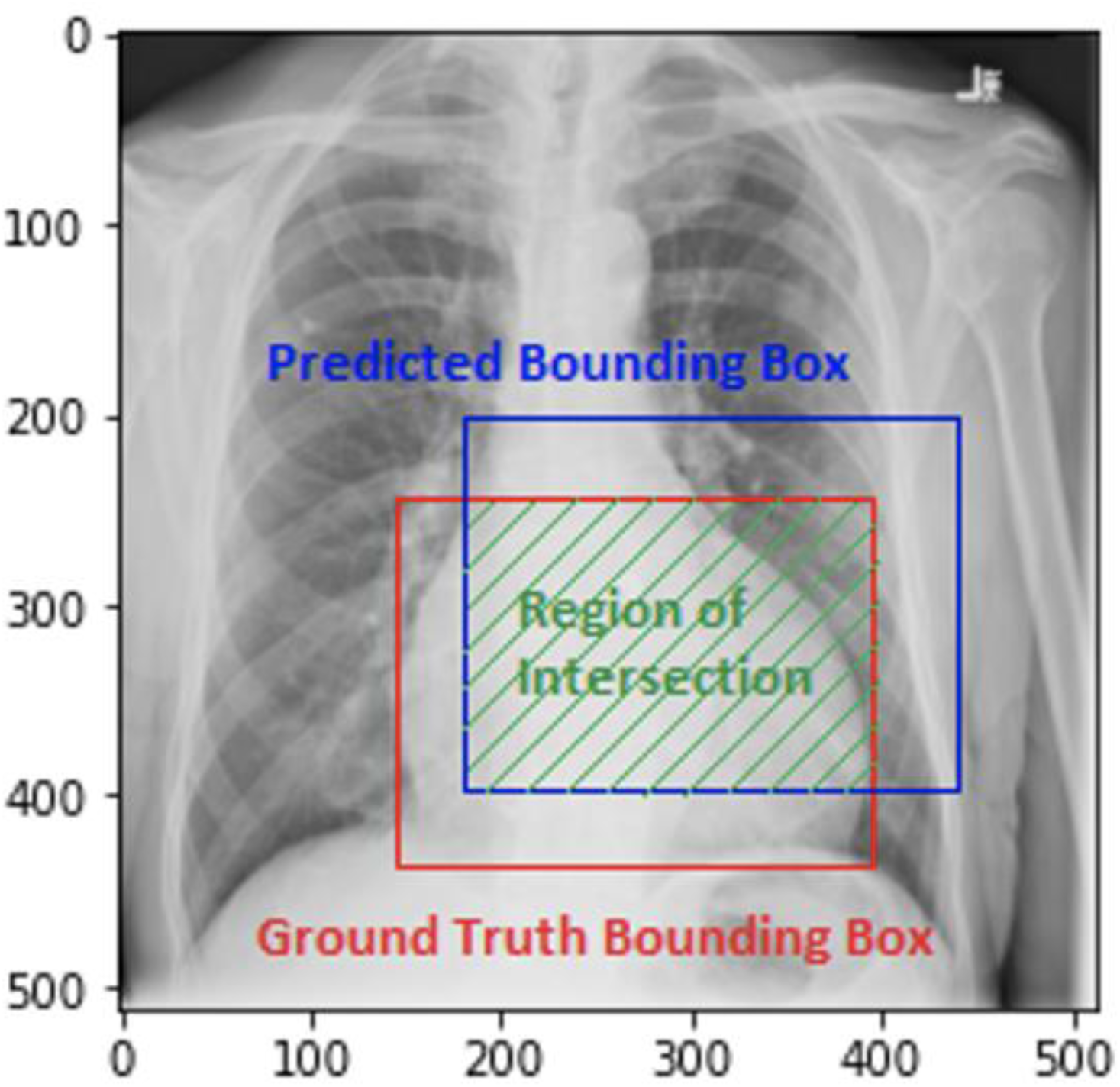
Image of a patient suffering from cardiomegaly showing intersection over union policy for the custom cost function.

## Results and Discussion

The predicted bounding box is found to provide an accurate position of the disease (cardiomegaly). The region of prediction is observed to be larger than that of the actual bounding box (figures 5 & 6). Although the region of bounding box was bigger than the actual ground truth bounding box, it seemed a reasonable offset. Further, training with additional images is likely to improve the box prediction score.

**Figure 6:**
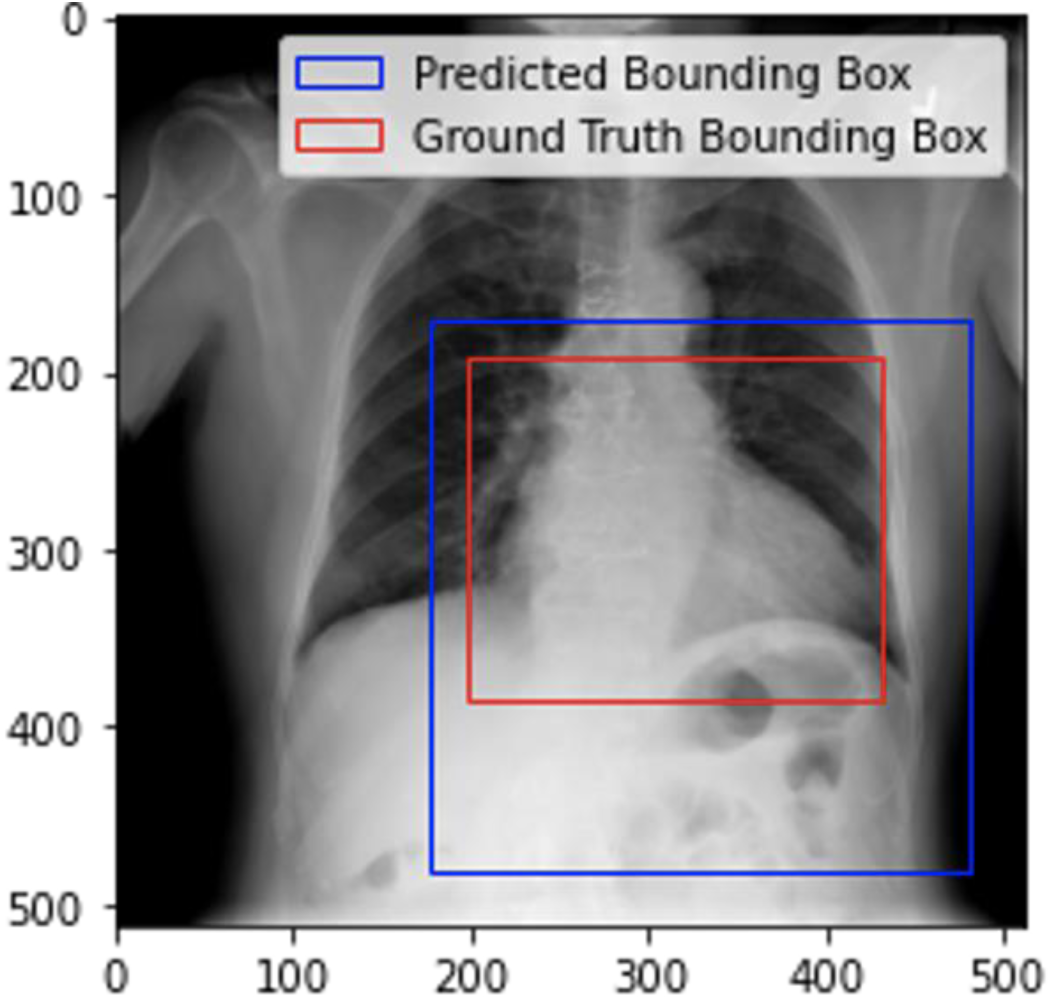
Comparison of ground truth bounding box with the bounding box predicted with SSD Model.

**Table 2:** Number of training epochs and losses for all algorithms.

| Bounding Box Algorithm | Optimizer Used | Learning Rate | No. of Training Epochs | Initial Loss | Final Loss |
| --- | --- | --- | --- | --- | --- |
| **Atelectasis** | Adam | 1e-4 | 50 | 11.91 | 1.23 |
| **Cardiomegaly** | Adam | 1e-2 | 38 | 2.95 | 1.05 |
| **Effusion** | Adam | 1e-3 | 50 | 7.85 | 1.09 |
| **Infiltration** | Adam | 1e-3 | 50 | 9.43 | 0.97 |
| **Mass** | Adam | 1e-3 | 50 | 10.64 | 1.19 |
| **Nodule** | Adam | 1e-3 | 50 | 8.24 | 1.50 |
| **Pneumonia** | Adam | 1e-3 | 50 | 7.45 | 0.89 |
| **Pneumothorax** | Adam | 1e-3 | 50 | 8.44 | 1.00 |

This seems a promising strategy of utilizing SSD with a VGG-16 network as a backbone for feature detection of bounding box algorithm to predict the location on X-ray images. Its applications should be tested on other medical image datasets like computerized tomography, magnetic resonance image or even immunohistochemistry staining images. Bounding boxes are one of the most popular image annotation techniques in deep learning, and with improvements in prediction accuracies, this method can reduce costs and increase annotation efficiency compared to other image processing methods.

## Supplementary data

1. National Institute of Health chest X-ray dataset: https://www.kaggle.com/nih-chest-xrays/data
2. Bounding box coordinates of the testing images: https://www.kaggle.com/nih-chest-xrays/data?select=BBox_List_2017.csv
3. Analysis code can be retrieved from here: https://bitbucket.org/chestai/chestai_rushikes_code/src/master/

## Author contributions

SP, AS, and RC conceived the concepts, planned, and designed the article. SP, AS, and RC primarily wrote and edited the manuscript.

